# Prenatal alcohol exposure alters expression of genes involved in cell adhesion, immune response, and toxin metabolism in adolescent rat hippocampus

**DOI:** 10.1101/2023.06.12.544533

**Authors:** Amal Khalifa, Rebecca A.S. Palu, Amy Perkins, Avery Volz

**Affiliations:** Department of Computer Science, Purdue University Fort Wayne, Fort Wayne, IN 46805; Department of Biological Sciences, Purdue University Fort Wayne, Fort Wayne, IN 46805; Department of Psychology, Purdue University Fort Wayne, Fort Wayne, IN 46805

## Abstract

Prenatal alcohol exposure can result in mild to severe consequences for children throughout their lives, with this range of symptoms referred to as Fetal Alcohol Spectrum Disorders (FASD). These consequences are thought to be linked to changes in gene expression and transcriptional programming in the brain, but the identity of those changes, and how they persist into adolescence are unclear. In this study, we isolated RNA from the hippocampus of adolescent rats exposed to ethanol during prenatal development and compared gene expression to controls. Exposure to ethanol caused widespread downregulation of many genes as compared to control rats. Gene ontology analysis demonstrated that affected pathways included cell adhesion, toxin metabolism, and immune responses. Interestingly, these differences were not strongly affected by sex. Furthermore, these changes were consistent when comparing ethanol-exposed rats to pair-fed controls provided with a liquid diet and those fed ad libitum on a standard chow diet. We conclude from this study that changes in genetic architecture and the resulting neuronal connectivity after prenatal exposure to alcohol continue through adolescent development. Further research into the consequences of specific gene expression changes on neural and behavioral changes will be vital to our understanding of the FASD spectrum of diseases.

**AUTHOR SUMMARY:** Alcohol exposure during fetal development is associated with a wide range of behavioral and physical symptoms that can be observed from childhood throughout adolescence and beyond. It is believed that this exposure may alter gene expression patterns permanently by changing genomic architecture, but the actual changes themselves are still unclear. In this study we examined gene expression patterns in rats whose mothers were given ethanol during their pregnancies. These were compared to control rats whose mothers were fed a similar liquid diet without ethanol as well as rats fed a normal diet. We identified the top differentially expressed genes and performed gene ontology analysis to identify both genes and pathways important in the response to ethanol during fetal development. We focused on adolescent rats since prenatal alcohol exposure has been shown at this stage to influence behavior. We indeed found a number of significant changes in gene expression, suggesting that prenatal alcohol exposure has ongoing consequences throughout and likely beyond adolescence into adulthood. Pathways such as cell adhesion, immune response, and toxin response were all highlighted. Future work will focus on making connections between these gene expression changes and behavioral changes observed at this same life stage.

## INTRODUCTION

Maternal consumption of alcohol is relatively common, with 10% of pregnant women reporting some drinking during pregnancy (Denny, Acero, Naimi, & Kim, 2019). Given that alcohol easily crosses the placental barrier, maternal alcohol consumption can significantly affect fetal development. Severe maternal alcohol consumption can result in Fetal Alcohol Syndrome, with milder consumption associated with Fetal Alcohol Spectrum Disorders (FASD). A conservative estimate of the prevalence of FASD in the United States is 1-5% (May et al., 2018). Prenatal alcohol exposure (PAE) is associated with a wide range of neurobehavioral outcomes, including altered sensory processing, intellectual disability, developmental delays, and cognitive dysfunction.

Neuroimaging has been critical in elucidating the neuroanatomical and functional consequences of PAE. Not only is PAE correlated with overall reductions brain volume and major fiber tracts (e.g., corpus callosum), but there are specific reductions in the volume of subcortical structures such as the hippocampus (Donald et al., 2015; Wozniak, Riley, & Charness, 2019). Functional MRI studies have reported altered neuronal activation during working memory and executive function tasks, but the data from neuroimaging studies has been inconsistent (Coles & Li, 2011). A recent study by Solar et al. (2022) used MRI and DTI to show that although the volume of the hippocampus was smaller in individuals with PAE, there were no significant group differences in mean diffusivity, axial diffusivity, or fractional anisotropy, suggesting that alcohol-related cognitive dysfunction is a result of changes at the cellular level (Solar, Treit, & Beaulieu, 2022). Rodent models have been critical in evaluating the short- and long-term effects of prenatal alcohol exposure at the neuronal level. For example, using unbiased stereology, Tran et al. (2003) found a significant reduction in neuron number in CA1 in adult rats following developmental alcohol exposure (Tran & Kelly, 2003). Furthermore, developmental alcohol exposure alters hippocampal synaptic plasticity (Valenzuela, Morton, Diaz, & Topper, 2012), which may help to explain alcohol-related cognitive deficits. Together, these studies point to the hippocampus as being a key site of alcohol-related alterations.

Using transcriptomics and high-throughput genetic screening techniques, researchers have demonstrated that prenatal alcohol exposure significantly alters the genome. For the most part, studies have shown that prenatal alcohol exposure results in a significant down-regulation of gene expression (R. Da Lee et al., 2004; D. H. Lee et al., 2016; Mishra, Shrinath, Rao, & Shukla, 2023; Rogic, Wong, & Pavlidis, 2016). In a study evaluating DNA methylation following prenatal alcohol exposure, buccal samples were obtained from newborns and evaluated for differentially methylated regions, with few such regions identified (Loke et al., 2021). However, others have looked at children and adolescents with FASD and reported significant differentially methylated regions. A study analyzing DNA methylation in buccal swabs (age range 3.5-18 years) reported a DNA methylation signature that may be unique to individuals with FASD (Lussier et al., 2018). DNA methylation alterations were also observed in blood samples in children and adolescents with FASD (Cobben et al., 2019). In addition, prenatal alcohol exposure alters gene expression in rodent fetal hippocampal tissue. Specific pathways identified as being altered by prenatal alcohol exposure include those involved in regulation of transcription (Mandal, Park, Lee, et al., 2015), G-protein coupled receptor signaling (Mandal, Park, Lee, et al., 2015; Mandal, Park, Jung, & Chai, 2015), proline and citrulline biosynthesis (Lunde-Young et al., 2019) and neuron differentiation (Mandal, Park, Jung, et al., 2015).

However, few studies have utilized genome-wide screens to evaluate the long-term consequences of prenatal alcohol exposure. Of those that have, a limited number have focused on adolescence as a key period of interest. In the amygdala (P42), prenatal alcohol exposure resulted in altered miRNA expression, with key pathways including those related to p53, CREB, glutamate and GABA (Ignacio, Mooney, & Middleton, 2014). Another study found that prenatal alcohol exposure resulted in significant upregulation of genes involved in neuroinflammation in the hippocampus of male adolescent rats (Marjonen et al., 2015). In the cortex, about 250 genes were downregulated as a result of prenatal alcohol exposure. Interestingly, far fewer genes were altered in female alcohol-exposed offspring relative to male alcohol-exposed offspring (37 vs. 271, respectively). Genes identified were associated with neurological disorders and the protocadherin family of proteins (Mishra et al., 2023). Finally, in a study comparing the effects of prenatal alcohol exposure on gene expression in the olfactory bulb of adolescent (P40) and adult (P90) rats, fewer changes were evident in adolescents. In fact, only a single gene was identified as being upregulated in ethanol-treated rats compared to pair-fed controls, dual specificity phosphatase 1 (*Dusp1*), whereas no genes were downregulated (Gano et al., 2020). Taken together, it is clear that prenatal alcohol exposure causes long-lasting alterations to the genome, but these effects vary by brain region. Furthermore, some studies have evaluated sex differences (Ignacio et al., 2014; Mishra et al., 2023), whereas others have utilized only male rats (Marjonen et al., 2015) or collapsed across sex (Gano et al., 2020). In the present study, we examined whether prenatal alcohol exposure resulted in changes in gene expression in the adolescent male and female hippocampus using RNA sequencing. We hypothesized that alcohol exposure would result in significant downregulation of gene expression, with highlighted genes involved in neuroinflammation, G-protein coupled receptor signaling, and cell adhesion.

## RESULTS

### RNA-sequencing differentiates ethanol-fed from pair-fed rats

To determine the impact of moderate prenatal ethanol exposure (BEC range: 14.90 - 110.70; mean: 44.03 ± 7.21) on gene expression in adolescent rats, RNA was isolated from the hippocampus of rats aged 36-38 days. Three dietary groups were compared: the offspring of females were either fed a normal chow diet ad libitum (AD), a liquid diet containing ethanol (ET), or a comparable liquid diet without ethanol (PF). Three males and three females were sampled from each cohort, with the exception of the alcohol-fed litter where only two females were available. Differential gene expression analysis was used to compare all pairwise combinations (adjusted p-value <0.01, Log2Fold-change>1.5), although particular attention was paid to ethanol compared to pair-fed samples since environmental variables related to diet were controlled, with the primary difference being presence or absence of ethanol in the maternal diet (Table S1-S6). Analysis identified 407 genes that were significantly downregulated upon prenatal exposure to ethanol, while 16 genes were upregulated as compared to PF rats (Table S1).

To determine whether biological sex has a significant impact on the expression of genes that respond to ethanol exposure, we compared expression profiles for males and females under each treatment condition as well as combined. In ET rats, 3 genes were significantly upregulated in males compared to females and 20 genes were upregulated in females compared to males (Table S5). In PF rats, 6 genes were significantly upregulated in males compared to females and 47 genes were significantly upregulated in females compared to males (Table S7). When these genes were collectively compared to those significantly altered in ET rats compared to PF rats, only one gene (*RT1-N2*) was found be affected by ethanol exposure (downregulated in ET rats) as well as differentially expressed between males and females (upregulated in females compared to males, only after exposure to ethanol). These results are supported by principle components analysis (PCA), which show that sex is not a primary factor segregating the samples (Figure S1). Considering this, we elected to continue with our analysis by pooling male and female samples under the same environmental conditions as conducted in previous studies (Gano et al., 2020).

PCA did show clustering of samples from the same environmental conditions (Figure 1A). The most distantly separated clusters do appear to be ET and PF groups, with AD samples appearing to bridge the gap between these two. This is consistent with previously published data, which has demonstrated differences in gene expression between PF and AD controls (Gano et al., 2020; Lussier et al., 2015). This clustering lends further support to the primary focus on PF and ET conditions, since these are not only the most directly comparable with regards to environmental conditions, but also appear to be the most differentiated in terms of gene expression.

**Figure 1.**
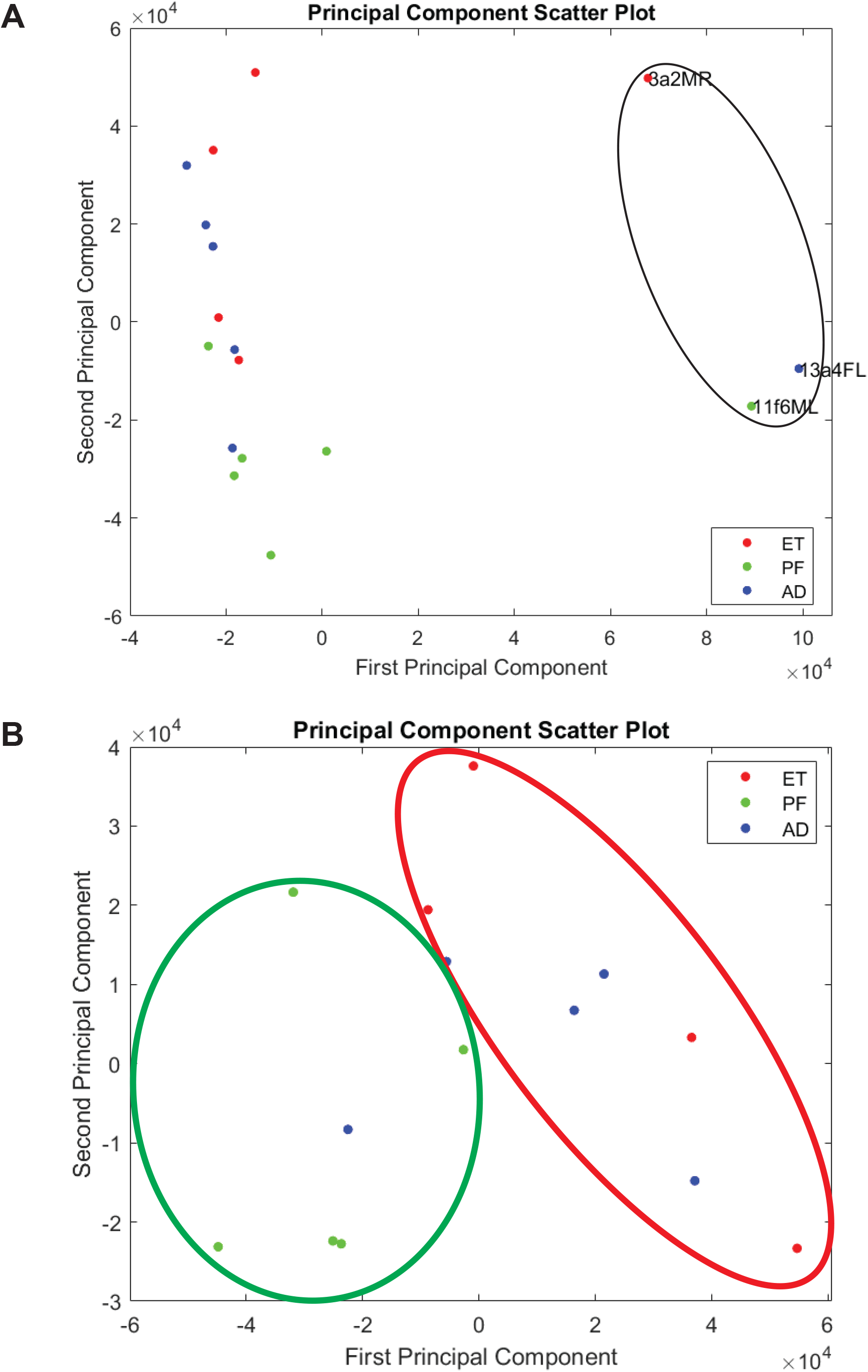
Feeding condition is a primary distinguishing factor in differential gene expression between ethanol-fed and pair-fed mice. **A.** Principle components analysis was performed on all 17 RNA-seq samples (first and second principle components). Ethanol exposure samples (ET) are marked in red, pair-fed control samples (PF) are marked in green, and ad libitum-fed control samples (AD) are marked in blue. ET and PF samples primarily segregate along the second principle component, with AD samples segregating between ET and PF. Three outlier samples, one from each group, segregate along the first principle component and are identified by a black circle. **B.** Principle components analysis was performed on all 14 RNA-seq samples after exclusion of the outlier samples marked above (first and second principle components). Ethanol exposure samples (ET) are marked in red (circled in red), pair-fed control samples (PF) are marked in green (circled in green), and ad libitum-fed control samples (AD) are marked in blue. ET and PF samples still segregate, now along both the first and second principle components, with AD samples segregating between ET and PF.

Additionally, PCA highlighted three potential outlier samples, one from each treatment group. These three samples were separated from the rest of the samples along the first principle component (Figure 1A). To ensure these outlier points did not have undue influence on the results, we removed them from all further analysis. 14 total samples (5 AD, 4 ET, and 5 PF) were compared in the final analysis, in which 393 genes were downregulated upon prenatal exposure to ethanol and 22 genes were upregulated. New PCA demonstrated that environmental condition was still a primary determinant of clustering even after removal of these samples (Figure 1B). Furthermore, 96% of these genes are shared with the original list, indicating that removing the outlier samples does not substantially change the overall results (Tables S7-S9).

### Genes involved in immunity and cell adhesion are altered after prenatal alcohol exposure

Differential gene expression analysis of the 14 samples remaining after filtering revealed 393 genes that were significantly downregulated upon prenatal exposure to ethanol compared to PF controls, while 22 genes were upregulated (Figures 2A, Table S7). Compared to the AD controls, 410 genes were significantly downregulated upon prenatal exposure to ethanol, while 79 genes were upregulated (Figure 2B, Table S8). Of these genes, 347 are downregulated in comparison to both PF and AD controls, while 16 are upregulated in comparison to both (Tables S7-S8). The overlap in effect is emphasized in a heat map for the top 10 up and down-regulated genes (ET vs. PF) with annotations (Figure 3). All ten most downregulated genes share a significant effect for ET compared to PF or AD. Seven of the ten most upregulated genes share a significant effect for ET compared to PF or AD. Taken together, these analyses suggest that the impact of ethanol feeding is consistent whether comparing to either PF or AD controls.

**Figure 2.**
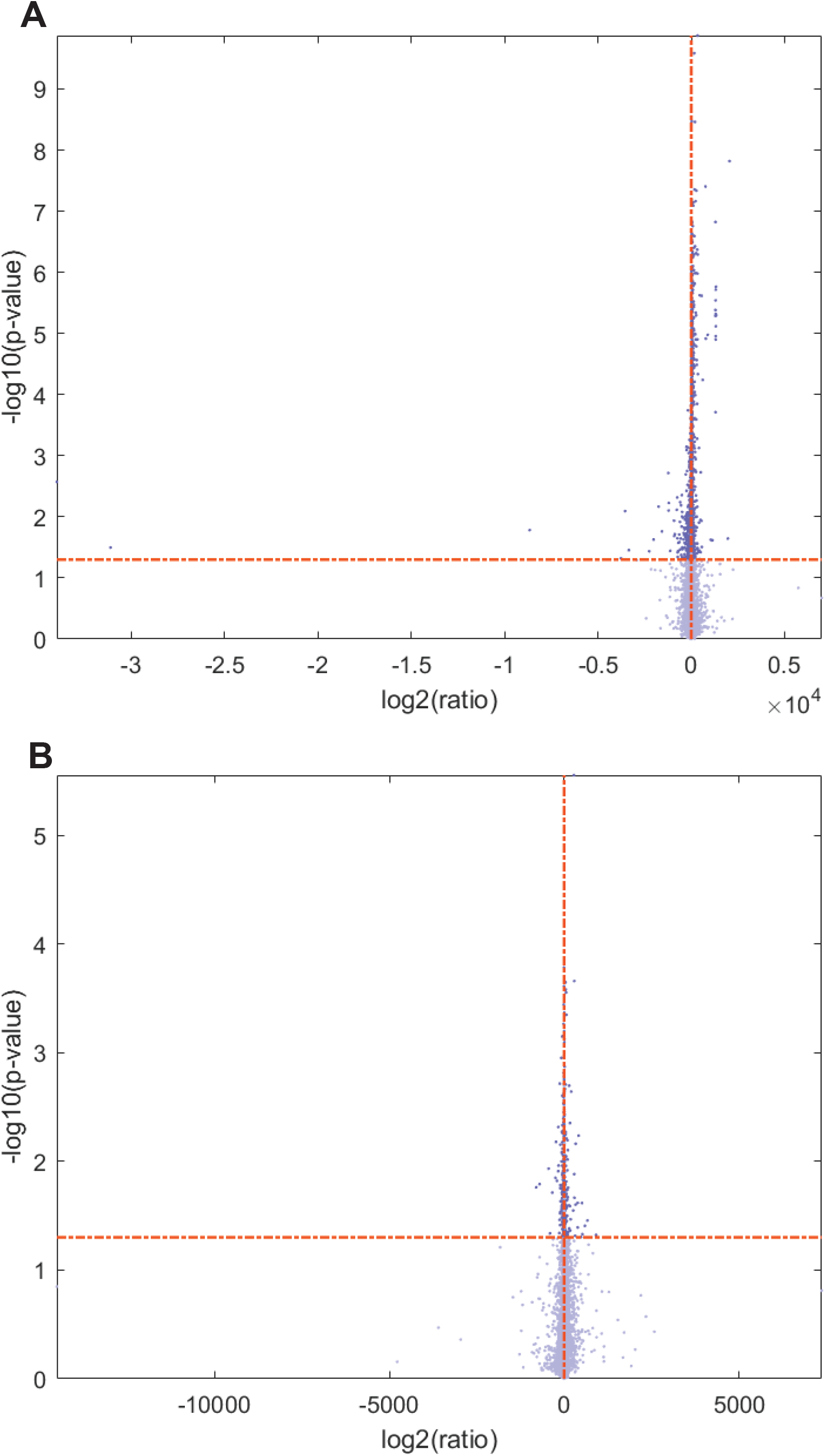
Ethanol exposure significantly alters expression of many genes as compared to pair-fed and ad libitum fed controls. **A.** A volcano plot was generated from data in Table S7 comparing ET rats to control PF rats. Log2 of the ratio of PF/ET is reported along the X-axis, and the log10 of the adjusted p-value for each change is reported along the Y-axis. Cut-offs for significance were set at an absolute value for Log2FC>1.5 and adjusted p-value of <0.01. These cut-offs are indicated as dotted lines on the plot. In total, 393 genes were significantly downregulated and 22 genes were significantly upregulated in ET rats compared to PF controls. **B.** A volcano plot was generated from data in Table S8 comparing ET rats to control PF rats. Log2 of the ratio of AD/ET is reported along the X-axis, and the log10 of the adjusted p-value for each change is reported along the Y-axis. Cut-offs for significance were set at an absolute value for Log2FC>1.5 and adjusted p-value of <0.01. These cut-offs are indicated as dotted lines on the plot. In total, 347 genes were significantly downregulated and 16 genes were significantly upregulated in ET rats compared to AD controls.

**Figure 3.**
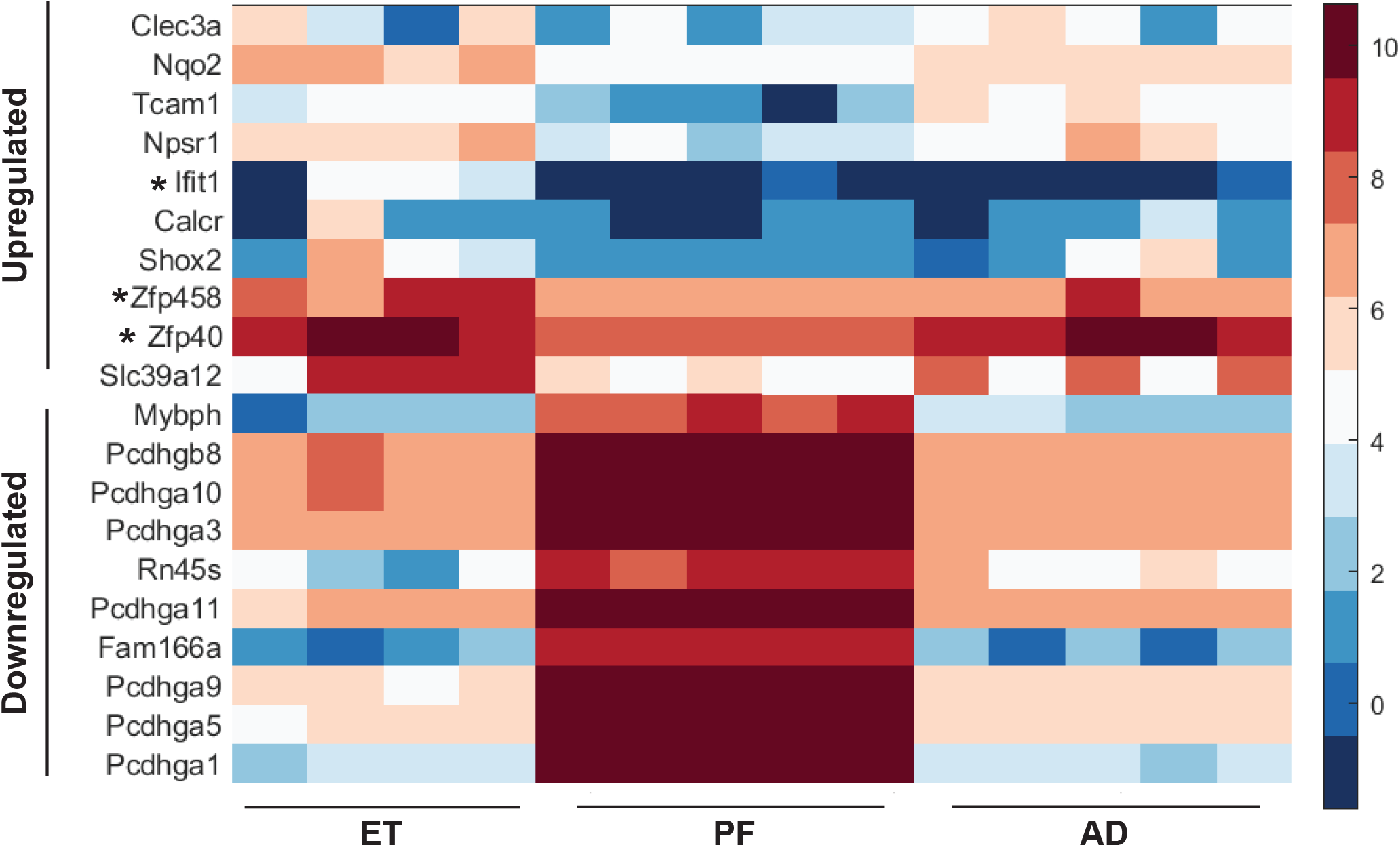
Top candidate genes differentially expressed in ET rats as compared to PF rats. A heat map was generated for the top 10 genes with annotations that were differentially expressed in ET rats compared to PF rats. Unannotated genes are defined as any with a location/annotation symbol only, and without an alternate gene name/function assigned. Individual samples for each group are shown as columns along the X-axis, with the expression of each gene in rows along the Y-axis. All ten downregulated genes are significantly changed in ET as compared to both PF and AD samples. Seven out of ten upregulated genes are significantly changed as compared to both PF and AD samples. Three genes (*Zfp40*, *Zfp458*, and *IFIT1*) are only significantly altered in ET as compared to PF samples (marked with an *). Both *IFIT1* and *ZFP458* show a trend toward increased expression in ET samples as compared to AD. Blue indicates low relative expression, while red indicates high relative expression (see key, log2(normalized read counts)).

Of the top ten most downregulated genes with functional annotation, seven belong to the protocadherin gamma subfamily A group of genes (Figure 3). The *PCDHGA* family of genes are encoded in a cluster and function in neuronal cell adhesion, particularly in the CNS, where they may play a role in the regulation of the blood-brain barrier (Dilling, Roewer, Fo, & Burek, 2017). This is complemented by other cell adhesion genes that are upregulated in response to ethanol exposure, such as *TCAM1*. The identification of cell adhesion in particular serves to validate our study, as previous work has demonstrated that prenatal alcohol exposure can alter cell-cell interactions through changes in cadherin expression during development (Licheri & Brigman, 2021). Changes in expression of these genes suggest that prenatal ethanol exposure may lead to changes in neuronal connectivity that persist well past fetal development into adolescence.

Another interesting gene found to be upregulated in ET rats as compared to PF rats is *IFIT1*, a conserved antiviral gene activated upon interferon signaling and commonly found to be upregulated in response to viral infection (Figure 3, Table S7) (Yan & Chen, 2012). This gene also appears to be upregulated in ET rats as compared to AD, although this change is not significant (Figure 3, Table S8). It is well known that alcohol exposure alters the neuroimmune response (Deak, Kelliher, Wojcik, & Gano, 2022). Thus, it is not surprising that our analysis identified several genes associated with immunity to be altered following prenatal alcohol exposure (Table S7).

### Cell adhesion, toxin metabolism, and immune processes are all downregulated after prenatal alcohol exposure

To determine if there are any pathways or processes that are significantly altered in ET rats as compared to PF rats when taking into account the entire gene list and not only the top candidates, we performed gene ontology analysis using the Database for Annotation, Visualization and Integrated Discovery (DAVID) (Huang, Sherman, & Lempicki, 2009; Sherman et al., 2022). Gene names were converted to Entrez gene IDs for analysis. *LOC100362040*, *LOC100366054*, and *LOC100363531* do not have current Entrez gene IDs and therefore were excluded from this analysis. No significant categories were identified in any analysis for genes upregulated in ET compared to PF rats (Table S10), likely due to the small number of significant genes identified.

We did observe, however a number of categories across analyses that were enriched in genes downregulated in ET versus PF rats (Table 1, Tables S11-S24). We pooled our results across multiple analyses of biological process, molecular function, pathway enrichment, and protein domains for further investigation (Table 1). Of particular interest was enrichment for cell adhesion and cadherin genes (IPR013164; SM00112, Cadherin, KW-0130, IPR020894, IPR002126, IPR015919, GO:7156, rno04514), glucuronidation (rno00040; GO:52696, GO:52697, GO:52695, UDPGT, IPR002213, rno00053, GO:15020), toxin metabolism (rno00983, rno00982, rno00980, rno05204, rno05207, rno04976), and the immune response (rno05332, rno05330, rno04940, rno05320, rno05416, rno04612) (Table 1, S11-S24). In all of these, multiple categories were identified through differing analyses, strengthening their association in this study. The further validation of cell adhesion as responding to prenatal alcohol exposure once more serves to validate our study (Licheri & Brigman, 2021). Our work supports these conclusions and furthermore indicates that they persist long after initial alcohol exposure, continuing into adolescence. Genes in these highlighted GO categories were compiled into a comprehensive list of 41 annotated genes (Figure 4). Of these 41 genes, 31 are also significantly downregulated in ET rats as compared to AD rats. Of particular note are the *PCDHGA* cell adhesion genes (12 of the 41 GO highlighted genes), as well as the *RT1 MHC* Class 1 genes (4 of the 41 GO highlighted genes) (Figure 4).

**Figure 4.**
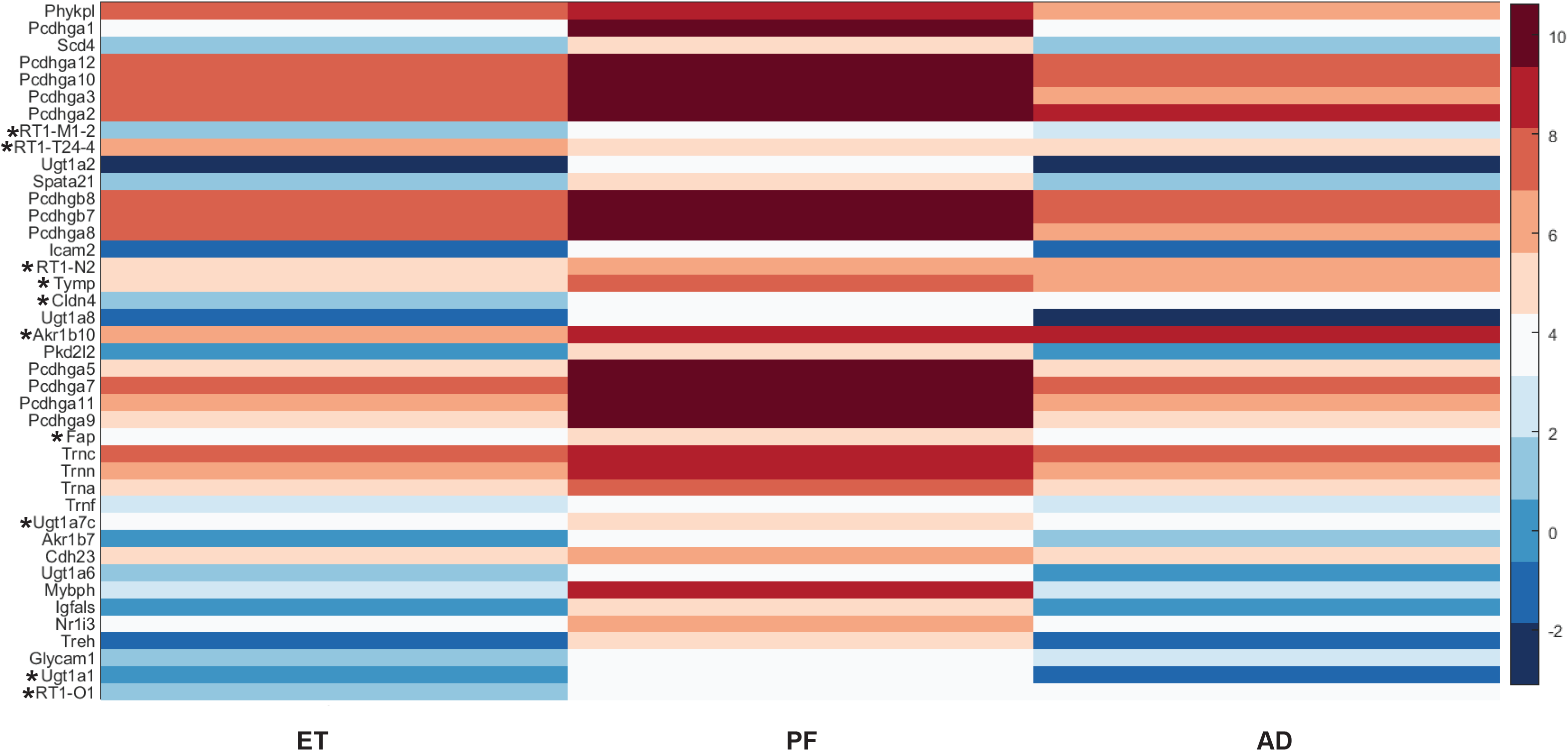
Genes from highlighted GO categories that are differentially expressed in ET rats as compared to PF rats. A heat map was generated for the 41 genes with annotations that were differentially expressed in ET rats compared to PF rats and were found in the highlighted GO categories from Table 1. Unannotated genes are defined as any with a location/annotation symbol only. Treatment categories are shown along the X-axis, with the average expression of each gene for that group in rows along the Y-axis. 31/41 genes are significantly downregulated in ET rats as compared to both PF and AD controls. Ten genes (*Akr1b10, Cldn4, Fap, RT1-M1-2, RT1-N2, RT1-O1, RT1-T24-4, Tymp, Ugt1a2, Ugt1a7*c) are only significantly altered in ET as compared to PF samples (marked with an *). In all except for *Fap* and *Ugt1a7c*, expression shows a trend toward decreased expression in ET samples as compared to AD. Blue indicates low relative expression, while red indicates high relative expression (see key, average log2(normalized read counts) across all individuals in a group).

**Table 1:** Processes enriched by Gene Ontology.

| Process of interest | GO Categories | Lowest P-value |
| --- | --- | --- |
| Cadherins and Cell Adesion | IPR013164; SM00112, Cadherin, KW-0130, IPR020894, IPR002126, IPR015919, GO:7156, rno04514 | 1.28E-14 |
| Glucuronidation | rno00040; GO:52696, GO:52697, GO:52695, UDPGT, IPR002213, rno00053, GO:15020 | 9.12E-09 |
| Toxin Response | rno00983, rno00982, rno00980, rno05204, rno05207, rno04976 | 4.25E-06 |
| Calcium ion binding | GO:5509 | 1.33E-05 |
| Misc Metabolic Processes | rno00860, KW-0328, rno00140, rno01240, rno00830, rno00970, rno01100 | 4.06E-05 |
| Immune Response | rno05332, rno05330, rno04940, rno05320, rno05416, rno04612 | 0.0026477 |

Glucuronidation was another interesting process highlighted by several analyses and categories (rno00040; GO:52696, GO:52697, GO:52695, UDPGT, IPR002213, rno00053, GO:15020). This pathway modifies various molecules with glucuronate, a sugar acid salt derived from glucose and most commonly involved in detoxifying and defense responses (Yang et al., 2017). During glucuronidation, the enzymes involved utilize UDP-glucuronate as a cofactor to add the sugar salt to the target. These molecules then are transported out of the cell, usually to be excreted in urine, bile, or feces. Common targets include toxins such as xenobiotics, drugs, and other environmental or dietary factors harmful to the organism (Yang et al., 2017). The downregulation of other toxin and drug metabolizing genes and pathways (rno00983, rno00982, rno00980, rno05204, rno05207, rno04976) in conjunction with glucuronidation suggests that the neurons continue to be affected by early toxin exposure in the form of alcohol.

In connection with this is the increased prevalence of several categories involved in the immune and defense response (rno05332, rno05330, rno04940, rno05320, rno05416, rno04612). Genes associated with different types of autoimmunity including graft-versus-host disease (rno05332), autoimmune thyroid disease (rno05320), and type I diabetes (rno04940) further indicates normal cellular defense mechanisms are downregulated in an attempt to adapt to prenatal alcohol exposure. Identification of *IFIT1* as a top upregulated candidate could indicate an attempt by the cells to counterbalance this general downregulation.

## DISCUSSION

Here we demonstrated that prenatal alcohol exposure produced persistent downregulation of gene expression in the hippocampus. Gene ontology analysis was used to determine the categories of genes that were affected by prenatal alcohol treatment. Several of these categories are of interest, with prenatal alcohol exposure reducing the expression of genes involved in cell adhesion, glucuronidation, toxin metabolism, and the immune response. In addition, few sex differences were evident, with only one gene (*RT1-N2)* upregulated in ET females compared to ET males. *RT1-N2* is predicted to play a role in antigen processing and is orthologous to human major histocompatibility complex class 1. Previous studies evaluating sex differences in gene expression in the CNS have had mixed results. For example, Gano et al. (2020) found no evidence of sex differences in the olfactory bulb of adolescent rats that were exposed to alcohol on GD11-12 (Gano et al., 2020). However, Mishra et al. (2023) reported that over 1,000 genes were differentially expressed in the cortex of male and female adolescent mice prenatally exposed to alcohol (Mishra et al., 2023). It is possible that sex differences in gene expression vary by brain region. Although most genes were downregulated following prenatal alcohol exposure, expression of *IFIT1* was upregulated in ET rats. *IFIT1* is part of the interferon-induced proteins with tetratricopeptide repeats (IFIT) family, where it participates in nonspecific antiviral response (Yan & Chen, 2012). It is somewhat surprising that prenatal alcohol exposure caused a long-lasting increase in the expression of *IFIT1* but supports other studies showing long-term programming of the immune response by alcohol (Deak et al., 2022; Gano et al., 2020).

We observed that prenatal alcohol exposure resulted in downregulation of several protocadherins (PCDH) from the gamma subfamily. PCDH are a subtype of cadherins, a superfamily of proteins involved in cell-cell interactions, that are expressed extensively in the central nervous system (Frank & Kemler, 2002). PCDH can be divided into several families: α, β, and γ, to name a few (Frank & Kemler, 2002). Of these, the γ-protocadherins (PCDH-γ) are of particular interest, due to their role in synaptogenesis (Weiner, Wang, Tapia, & Sanes, 2005) and blood-brain barrier formation (Dilling et al., 2017). Previous studies have found that prenatal alcohol exposure does not affect the expression of synaptophysin, PSD-95, or hippocampal spine density at PD30 (Elibol-Can, Kilic, Yuruker, & Jakubowska-Dogru, 2014; Jakubowska-Dogru, Elibol, Dursun, & Yürüker, 2017) or at PD60 (Jakubowska-Dogru et al., 2017). Thus, it is not likely that prenatal alcohol exposure results in long-term effects on synapse formation. However, PCDH-γ are expressed in endothelial cells of the blood-brain barrier (Dilling et al., 2017) and in astrocytes (Garrett & Weiner, 2009). Previous studies have reported that prenatal alcohol exposure disrupts brain vasculature in the hippocampus (Kelly, Mahoney, & West, 1990) and cortex (Siqueira, Araujo, Gomes, & Stipursky, 2021). Thus, it is possible that prenatal alcohol exposure disrupts blood-brain barrier integrity through a decrease in the expression of *PCDH-γ*. Although we did not evaluate gene expression in different cell types, it would be interesting to examine *PCDH-γ* expression in astrocytes, specifically. Astrocytes contribute to the formation of the blood brian barrier, and PCDH-γ are important for neuron-astrocyte interactions (Garrett & Weiner, 2009).

We observed significant differences in the expression of several genes involved in the immune response. First of all, several members of the *RT1* complex (rat MHC) class 1 family of genes were downregulated in the hippocampus of ET rats compared to PF rats *(RT1-M1-2, RT1-O1, RT1-N2, RT1-T24-4).* The *RT1* complex is a large cluster of genes, some of which are orthologous to human major histocompatibility complex (MHC). In the rat, the *RT1* complex can be further divided into framework, Class 1, and Class II genes (Dressel, Walter, & Günther, 2001; Günther & Walter, 2001). MHC class 1 molecules and their receptors are expressed in many neuronal subtypes as well as in microglia, where they are thought to play a pivotal role in the refinement of synapses (Cullheim & Thams, 2007). Thus, the observed downregulation of *RT1* class 1 genes following prenatal alcohol exposure suggests that important developmental processes, such as synaptic pruning and stripping, may be impaired by PAE.

Another important component of the immune response involves neutrophil transfer across the blood-brain barrier, a process that relies on intercellular cell adhesion molecule (ICAM)-1 and ICAM-2 (Lyck & Enzmann, 2015). ICAM-2 is expressed on endothelial cells (de Fougerolles, Stacker, Schwarting, & Springer, 1991; Fabry et al., 1992) and plays a role in T-cell diapedesis through the blood-brain barrier (Steiner et al., 2010). Furthermore, an analysis of fetal expression of *ICAM-2* revealed colocalization with microglia in the corpus callosum and cerebral vessels of the cortex (Rezaie & Male, 1999), although this study observed low expression of *ICAM-2* in the adult brain. However, a study by Navratil et al. (2003) reported significant expression of *ICAM-2* in vascular endothelial cells of adult post-mortem tissue (Navratil, Couvelard, Reyt, Hénin, & Scoazec, 1997). We observed a significant decrease in *ICAM-2* following prenatal alcohol exposure, which when combined with the observed downregulation of *PCDH-γ* could indicate significant dysfunction of the blood-brain barrier in alcohol-exposed rats. A previous study found that prenatal alcohol exposure resulted in altered angiogenesis and blood-brain barrier composition in newborn mice (Siqueira et al., 2021), whether these changes persist into adolescence remains to be seen. The data presented here suggests that prenatal alcohol exposure could indeed result in long-term changes to the integrity of the blood-brain barrier.

Taken together, it is clear that prenatal alcohol exposure causes enduring changes to the hippocampal genome. Almost 400 genes were downregulated in ET rats compared to PF rats, with GO analysis indicating enrichment of cell adhesion and cadherin genes, genes involved in toxin metabolism, and genes with a role in the immune response. Of note is the observation that several *PCDH-γ* and *RT1* complex genes were downregulated by alcohol exposure. Given the role of these genes in synapse formation and synaptic refinement, it is possible that prenatal alcohol exposure impairs critical processes such as synaptic pruning within the hippocampus. The hippocampus plays a major role in synaptic plasticity and memory consolidation, so impaired synaptic pruning within this structure could contribute to cognitive dysfunction in individuals with FASD.

## MATERIALS AND METHODS

### Subjects

Adult male and female Long-Evans rats were purchased from Charles River Laboratories (Wilmington, MA). Male and female pairs were co-housed overnight until a sperm plug was observed. At this time, females were removed and single-housed. Dams were assigned to one of three treatment groups: Ad-libitum control (AD), Pair-fed control (PF), and Ethanol-exposed (ET). Ad-libitum control dams were given *ad libitum* access to standard rat chow and were weighed daily throughout gestation. Pair-fed control rats received a control liquid diet (Dyets, Bethlehem, PA; Weinberg-Keiver Heigh Protein Control Diet 710324) and Ethanol-exposed rats received an experimental liquid diet (Dyets, Bethlehem, PA; Weinberg-Keiver Heigh Protein Experimental Diet 710109) with 36% ethanol-derived calories. PF dams were yoked to an ET dam of similar weight, such that the PF dam received the same amount of control diet, adjusted for body weight. To facilitate consumption of the ethanol liquid diet, ET dams were gradually exposed to the diet (Day 1: 33% experimental diet, 67% control diet, Day 2: 67% experimental diet, 33% control diet, Day 3: 100% experimental diet). Fresh diet was provided daily within 1.5-2 hours of lights off (Holman, Ellis, Morgan, & Weinberg, 2018). All dams were provided water ad libitum. On Gestational day (GD) 15, a small tail (∼ 1 mL) blood sample was taken from a separate set of ET rats (n = 17) to determine blood ethanol concentrations (BECs). Samples were also taken from PF rats to control for the stress of the tail blood procedure. The average (± SEM) BEC from this group of dams was 44.03 ± 7.21 (Range = 14.90 - 110.70). On GD 20, all rats were switched to standard food and water.

Body weights were collected from all offspring on postnatal day (PD) 7, 14, 21, and 35-37. Overall, there were few differences in body weights, with the exception that males weighed significantly more than females on P7 (p < 0.05). No effects of sex or condition were evident at PD14 or PD21. Male and female offspring were housed with their dam until weaning on PD21. After weaning, rats were housed in same-sex groups of 2-3 rats per cage until tissue collection. There were no significant effects of condition on body weights at the time of tissue collection, although males weighed significantly more than females [F(1,17) = 53.12, p < 0.0001]. Overall, these data were consistent with expectations, with males weighing more than females, and no effect of prenatal ethanol exposure.

### Hippocampus tissue collection

Rats were 36-38 days of age at time of sample collection. Male and female rats from each treatment group (n = 2-3/group) were removed from their home cage and rapidly decapitated without anesthesia. Whole brains were removed and hippocampi were dissected on ice, flash frozen, and stored at -80°C until RNA isolation.

### RNA isolation

RNA was isolated from frozen hippocampal samples stored at -80°C using the Monarch® RNA Total Isolation kit (NEB T2010S). Hippocampus from the left hemisphere was used where possible, with right hippocampus used for 4 samples. Samples were homogenized frozen in 1X DNA/RNA Protection Reagent (NEB T2011L), then processed according to kit instructions including on-column DNase digestion. Samples were eluted in 50 µL of RNAse-free water and frozen at -80°C until submitted for sequencing.

### RNA-sequencing

Concentration and purity of RNA was determined using a Nanodrop One C (ThermoFisher) prior to sequencing. Library generation (Illumina TruSeq Stranded mRNA HT Library) and sequencing (NextSeq 75 – High Output) were performed by the Center for Genomics and Bioinformatics at Indiana University.

### Quality Control and Genome Mapping

The sequenced data consists of a total of 51 paired-end sequence reads over three runs (replicates). This raw read data is saved in FASTQ-format files where each sequence has approximately 22 million reads. Those reads were pre-processed using Fastp (version 0.23.2); an all-in-one multithreaded high-performance Quality control tool (Chen, Zhou, Chen, & Gu, 2018). We employed the default parameters in Fastp where all sequence adapters were auto detected and trimmed. The reads are scanned with the sliding window of 4 and low-quality bases are filtered if the quality per base drops below 15 with 40% of bases are allowed to be unqualified. The minimum length to detect polyG in the read tail is 10. Furthermore, reads shorter than 15 or those with the number of N base is greater than 5 will be discarded.

The resulted filtered reads (approximately 93%) were mapped against the Rattus norvegicus (mRatBN7.2) genome assembly. The read mapping process was performed using Bowtie2 (Version 2.4.2) on the Galaxy platform (https://usegalaxy.org/; version 2.0.1) using the default options (Langmead & Salzberg, 2012). Next, the FeatureCounts tool (part of the Subread package version 2.0.1) was used to quantify the number of reads mapping to the exons of each gene (Liao, Smyth, & Shi, 2014). For a total number of 34322 genes, fragments (or templates) were counted instead of reads. The minimum mapping quality per read was 10 and reads were allowed to map to multiple features. The output of FeatureCounts is a table that follows the simplified annotation format (SAF) containing counted number of reads (fragments) mapped to each gene (Entrez gene identifiers).

### Differential Gene Expression Analysis

The primary objective of these experiments is to detect transcripts showing differential expressions across various conditions. Among the statistical approaches that have been designed to solve this problem, the DESeq method has been proven to be efficient in finding genes that are differentially expressed between two conditions (Soneson & Delorenzi, 2013). It is based on a negative binomial distribution model which has been shown to best fit the distribution of read counts across biological replicates (Anders & Huber, 2010).

In this research, differential analysis was performed using the rnaseqde() function that is implemented as part of the *Bioinformatics Toolbox* in *Matlab* (R2022a). As it follows the DESeq method, the rnaseqde() function estimates the biological variance between two conditions using at least two replicates for each. The count data is usually stored as a big table with gene identifiers in the rows and samples in the columns. Prior to performing the hypothesis test, the function applies a normalization step (median-of-ratios) to account for differences in sequencing depth sequencing depth and library composition between samples.

The output of rnaseqde() function is a summary table that includes (for each gene) the mean normalized counts averaged over all samples from both conditions, the fold change in log2, the P-value from the hypothesis test, and the adjusted p-value calculated using a False discovery rate (FDR) method. By default, the function uses the linear step-up procedure introduced by Benjamini and Hochberg. In this study, genes that satisfied FDR adjusted P-value less than 0.01 and an absolute fold change above 1.5 were considered differentially expressed. This cut off will identify the most up-regulated or down-regulated genes with at least a 1.5-fold change.

### Modifier Gene Characterization

Information about candidate genes and their human orthologues was gathered from a number of databases including Ratmine, OMIM, and NCBI, then verified through primary sources. Gene ontology categories were derived using the Database for Annotation, Visualization and Integrated Discovery (DAVID) (Huang et al., 2009; Sherman et al., 2022). Gene names and annotation symbols from RNA-seq were converted where possible to Entrez GeneIDs for analysis. This was done using DAVID for most genes, and the rest were identified individually through NCBI Gene (https://www.ncbi.nlm.nih.gov/gene/). Three genes (*LOC100362040*, *LOC100363531*, and *LOC100366054*) did not have current Entrez GeneIDs and were therefore excluded from analysis. Results from default analyses are included in Tables S10-S24. Categories were considered significant if p > 0.05 after Benjamini multiple testing correction.

### Ethics Statement

All animal procedures were approved by the Institutional Animal Care and Use Committee at Purdue University (Protocol #1912001989).

## ACKNOWLEDGEMENTS

We would like to thank Carolyn Pang and Jayde Bransteter for their contributions to this project. Funding provided by Purdue University Fort Wayne through a Collaborative Research Grant from Purdue University Fort Wayne to AK, RASP, and AP. Support was also provided to RASP and AP from the Purdue University Fort Wayne Departments of Biology and Psychology.

## SUPPLEMENTAL INFORMATION

**Supplemental Figure 1. Sex is not a primary factor in differential gene expression among samples.** Principle components analysis was performed on all 17 RNA-seq samples. The first principle component segregates along the X-axis, and the second principle component segregates along the Y-axis. Male samples are marked with blue, while female samples are marked with red. Sex does not appear to be a primary determinant in the distribution of samples along either the first or second principle component.

**Supplemental Figure 2. Genes from highlighted GO categories that are differentially expressed in ET rats as compared to PF rats, broken down by sample.** A heat map was generated for the 41 genes with annotations that were differentially expressed in ET rats compared to PF rats and were found in the highlighted GO categories from Table 1. Unannotated genes are defined as any with a location/annotation symbol only. Individual samples for each group are shown as columns along the X-axis, with the expression of each gene in rows along the Y-axis. 31/41 genes are significantly downregulated in ET rats as compared to both PF and AD controls. Ten genes (*Akr1b10, Cldn4, Fap, RT1-M1-2, RT1-N2, RT1-O1, RT1-T24-4, Tymp, Ugt1a2, Ugt1a7*c) are only significantly altered in ET as compared to PF samples (marked with an *). In all except for *Fap* and *Ugt1a7c*, expression shows a trend toward decreased expression in ET samples as compared to AD. Blue indicates low relative expression, while red indicates high relative expression (see key, log2(normalized read counts)).

